## Supplementary Figure for "CoVET: Cryo-Electron Tomography guided by optical electrophysiology"

**LIST OF SUPPLEMENTAL INFORMATION**

**Supplementary Fig. 1.** Voltage imaging of HEK293T cell with electric field stimulation

**Supplementary Fig. 2.** Overall workflow of subtomogram analysis of ribosomes

**Supplementary Fig. 3.** FSC curve of the consensus map and each conformation

**Supplementary Fig. 4.** Statistical analysis of each conformational state of polysomes in different clusters

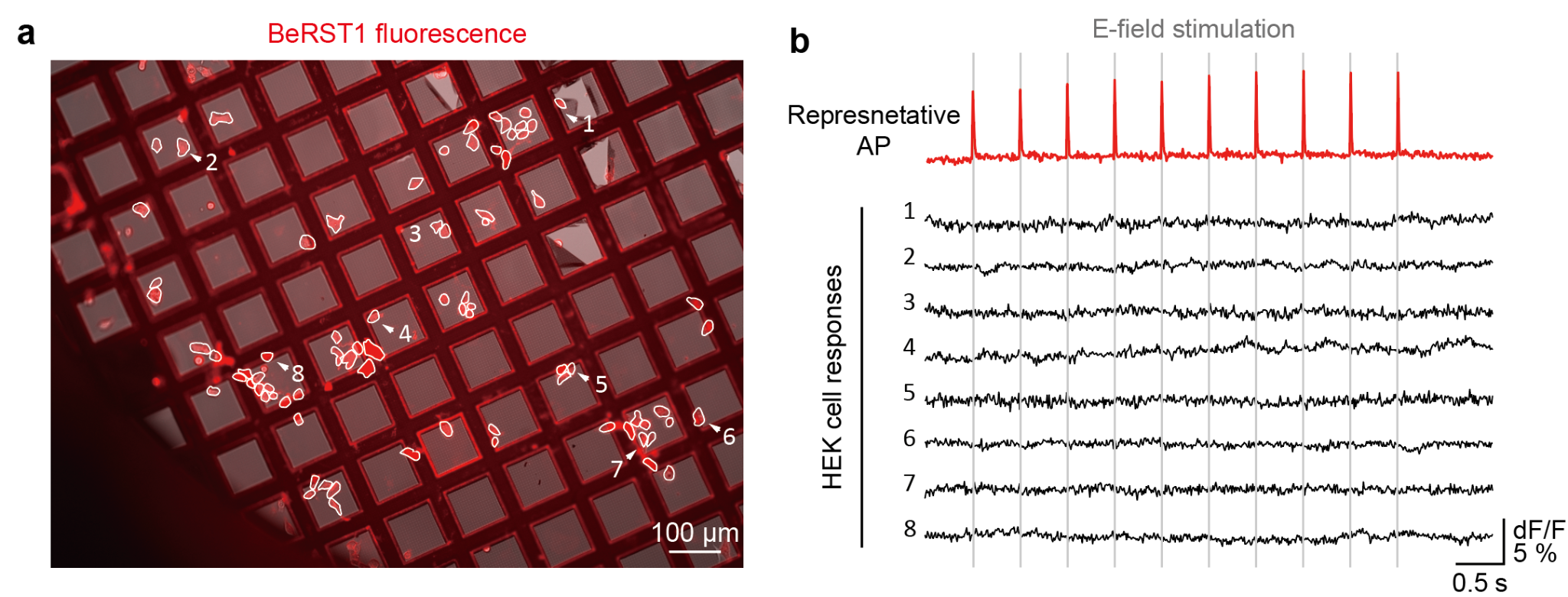

**Supplementary Fig. 1**. Voltage imaging of HEK293T cell with electric field stimulation

**a.** On-grid voltage imaging of HEK293T cells during electric field stimulation. HEK293T cells on a grid were stained with the dye BeRST1 (red). **b.** Representative Δ*F/F* fluorescence intensity changes according to time from annotated HEK293T cells in **a**. The shaded line indicates the electric field stimulation time.

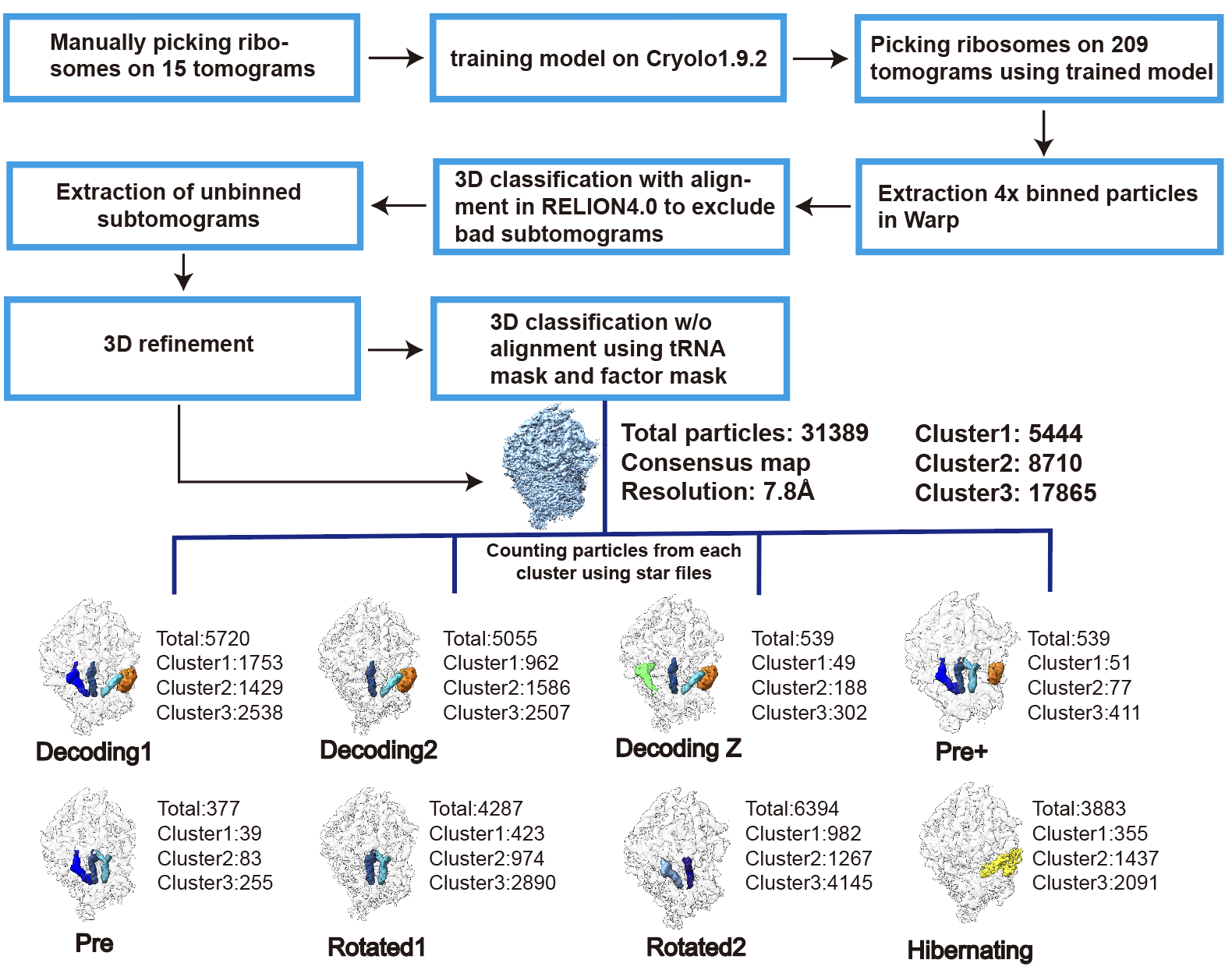

**Supplementary Fig. 2**. Schematic diagrams of the overall workflow. Deep learning-based picked subtomograms were extracted and filtered using iterative 3D classification with alignments. Re-extracted 2x binned subtomograms and masks, including the E (Exit), P (Peptidyl), and A (Aminoacyl) sites, along with elongation factors, were used to classify 8 different ribosome classes. Information from Starfiles was utilized to count the number of particles in each state from different clusters, which were then used for statistical analysis.

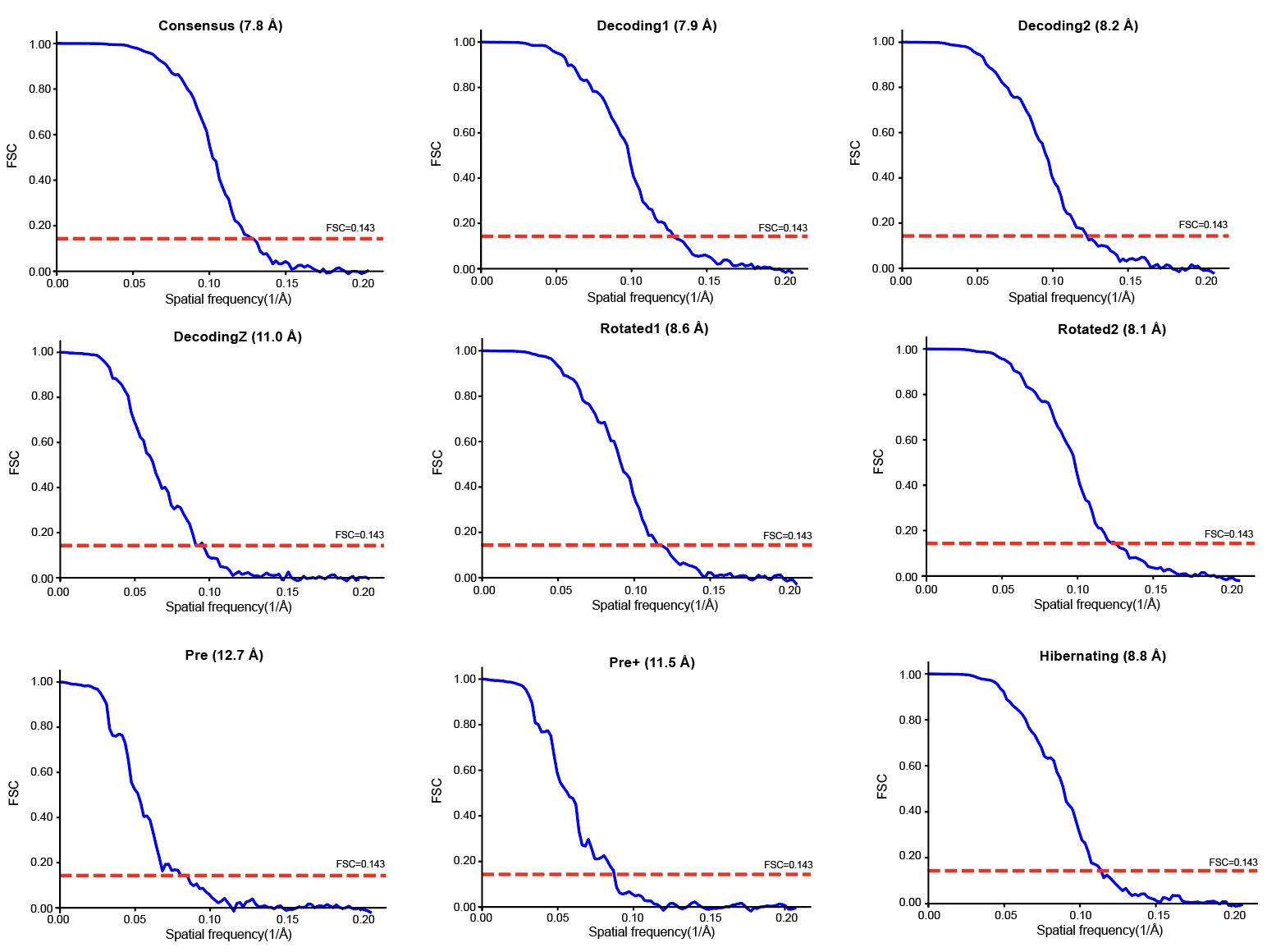
**Supplementary Fig. 3**. FSC curve of the consensus map and each conformation

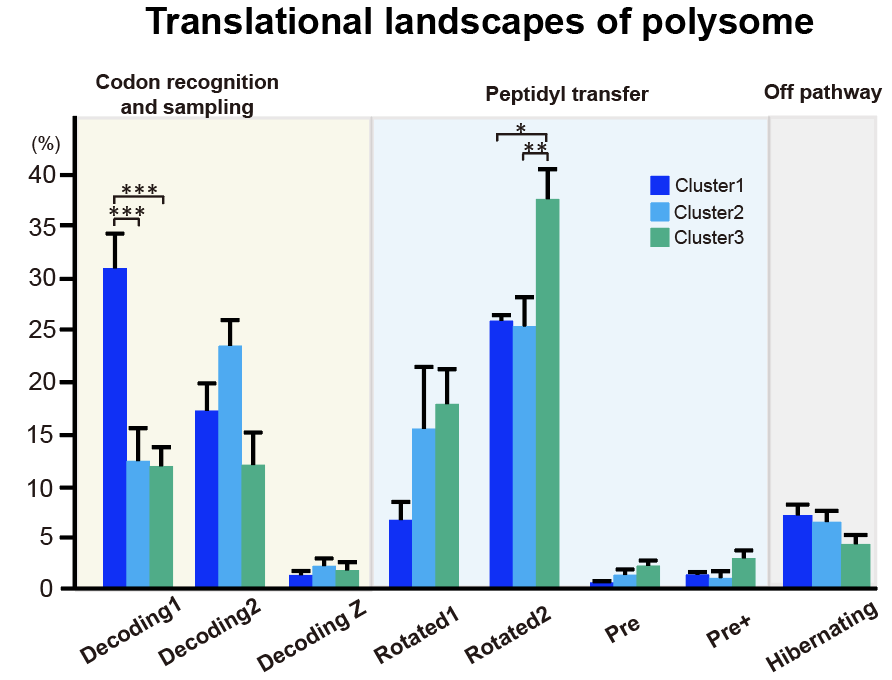

**Supplementary Fig. 4**. Statistical analysis of each conformational state of polysomes in different clusters

The conformational states of the polysomes where the distance between ribosomes was less than 9 nm were plotted. One-way analysis of variance (ANOVA) followed by Tukey’s post hoc test for multiple comparisons was performed to assess statistical significance. * *P* < 0.05, ** *P* < 0.01, *** *P* < 0.001.
